# ddkg.skill: A Compositional Agent Skill for Translating Biomedical and Bioinformatics Questions into Cypher for the Data Distillery Knowledge Graph

**DOI:** 10.64898/2026.09.26.754400

**Authors:** Deanne M. Taylor, Aditya Lahiri, Benjamin Stear, Taha Mohseni Ahooyi, Yuanchao Zhang, Asif Chinwalla, Shiping Zhang, Christopher Nemarich, J. Alan Simmons, Jonathan C. Silverstein

## Abstract

Biomedical knowledge graphs can connect information across genes, phenotypes, tissues, pathways, experiments, and clinical resources, but they are difficult to query correctly without detailed knowledge of the graph. Large language models can help write Cypher code for knowledge graphs, yet a query focused on a bioinformatics task that looks reasonable may still use the wrong identifier, relationship direction, source, intermediate node, or output unit if the user is not completely trained on the schema. We developed ddkg.skill, an Agent Skill that can be loaded into a compatible LLM session to support querying of the NIH Common Fund Data Ecosystem Data Distillery Knowledge Graph (DDKG), a biomedical property graph integrating more than 180 ontologies, over 40 genomics datasets, and dozens of additional cross-domain biomedical datasets. Rather than utilizing a simple markdown prompt, ddkg.skill contains a controller, a compositional library of DDKG-specific references and structured tables, primary documentation, validated query patterns, a routing table, and a Python script that checks the internal links among these materials. The skill build for the December 2025 DDKG release contains 38 bundled files and 239 routing relationships and is identified by checksum so users can state the exact build used to compose a query. The evaluation reported here was performed separately on an earlier build with 38 files and 227 routing entries. In that orthogonal nine-test evaluation against a live December 2025 DDKG instance, five of seven tests targeting sources absent from the worked examples produced correct executed results. The evaluation also exposed a failed query, a cross-source comparison that was not biologically well posed, and errors in the skill’s own reference material that informed later revisions. The skill guides an AI through entity resolution, graph inspection, query construction, and validation for biomedical and bioinformatics queries, while keeping the DDKG itself as the source of returned results. This design provides a portable and versioned method for giving general-purpose AI systems practical knowledge of a complex biomedical knowledge graph to empower complex bioinformatics data integration. Three executed biomedical use cases further demonstrate cross-species phenotype-to-expression querying, genomic overlap between 4DN chromatin-loop anchors and GTEx eQTLs, and integration of congenital heart phenotype associations with ClinVar and GTEx heart evidence.

## Introduction

Biomedical knowledge graphs (KGs) bring information from otherwise separate databases and ontologies into a form that can be queried across sources. The Data Distillery Knowledge Graph (DDKG) is built on the Unified Biomedical Knowledge Graph (UBKG) framework and was developed off our earlier Petagraph work [1]. UBKG utilizes the Unified Medical Language System (UMLS) [2] with additional biomedical ontologies and standards to connect source-specific biomedical codes and terms. DDKG is a large, heterogeneous biomedical property graph that builds on this framework by integrating genomics and supporting data from NIH Common Fund resources such as GTEx, HuBMAP, LINCS, 4DN, Kids First, and MoTrPAC with widely used biomedical resources including ClinVar, ClinGen, the Human Phenotype Ontology, Reactome, GENCODE/HGNC, STRING, and UniProt among several others [3,4]. This framework allows DDKG to support questions that cross genomic features, genes and proteins, tissues and cell types, phenotypes, pathways, clinical variables and experimental datasets. Other open-source approaches to biomedical KGs, such as the PheKnowLator ecosystem, use different graph models and construction strategies [5]. DDKG also incorporates resources developed specifically to support cross-domain queries, including the Homo Sapiens Chromosomal Location Ontology, which provides a common representation of genomic location and can connect datasets across genomic proximity [6].

Despite its demonstrated utility, a practical problem is that its breadth makes DDKG difficult to casually query without detailed knowledge of its schema. A query writer needs to be familiar with the community identifiers, data sources, relationships, and DDKG graph structures used to represent the interrogated biology. Knowing Cypher syntax alone is not sufficient without understanding the schema or the biological question being asked. A syntactically valid query can use the wrong source or relationship, miss an intermediate structure, count the same biological entity more than once, or return an empty result even without producing a database error.

Several existing approaches point to solutions to these challenges using modern artificial intelligence (AI). CypherBench and Mind the Query evaluate the use of large language models (LLMs) with KGs to translate natural-language questions into Cypher, and show that large schemas and execution against real KGs still remain challenging [7,8]. In biomedicine, BioChatter combines LLMs with graph-specific information to support natural-language interaction with BioCypher knowledge graphs [9,10]. The successful use of LLMs to help users navigate complex biomedical queries is encouraging, but generating valid KG queries is only part of what is required for an LLM to help navigate complex biomedical schemas like DDKG’s. An expert DDKG user could instead provide an LLM with explicit knowledge of how a particular DDKG release represents individual biomedical resources, which identifiers and relationships are appropriate for a given user question, and which query constructions can produce incomplete or misleading results. The LLM could then be used to assist other users in exploring the DDKG by helping generate appropriate queries on demand.

A complementary approach is to expose knowledge-graph services directly to AI systems through the Model Context Protocol (MCP). The NSF Prototype Open Knowledge Network has developed an MCP service for the Open Knowledge Network (OKN.us) that gives AI assistants programmatic access to a federation of scientific knowledge graphs. The service can discover relevant graphs, inspect their schemas, execute SPARQL queries, bridge identifiers between graphs, and combine results from multiple graph sources. This architecture places graph discovery and execution behind a standardized tool interface, allowing an MCP-capable LLM to query live knowledge resources rather than relying only on information supplied in its conversational context [16]. However, a limitation of the MCP concept is that querying of graphs can only be executed within the context for which the MCP server was designed.

Agent Skills provide another versatile mechanism for supplying task-specific knowledge to general-purpose AI systems. Originally introduced by Anthropic and subsequently released as an open standard, Agent Skills are a lightweight format adopted by major LLM vendors for packaging reusable instructions, scripts, reference materials, and other resources that an AI agent can discover and load when relevant to a task [11]. Rather than modifying model weights, this approach addresses the gap between a model’s general capabilities and the procedural, organizational, or domain-specific knowledge needed to perform specialized work reliably. Skills can therefore be versioned, shared, and reused across compatible agent environments while allowing detailed supporting material to be loaded only when needed. For knowledge-graph applications, a skill can encode the specialized procedural and release-specific knowledge needed to use a graph appropriately without itself providing a live interface to that graph. Applying an Agent Skill to DDKG therefore raises the possibility that the knowledge required to query a particular biomedical graph, including its identifiers, source conventions, relationships, and known failure modes, can itself be packaged as a versioned resource for a general-purpose AI system rather than learned through training a new text-to-Cypher model. An advantage of the Agent Skill is that it is natively flexible in the LLM context; it is not limited to a certain set of queries, unless the skill is specifically designed to be restrictive.

Evidence for the overall benefit of using Agent Skills remains mixed, although two recent evaluations are more consistent than they first appear. SkillsBench v1 evaluated 86 tasks across 11 domains under matched conditions and reported that curated skills increased average pass rate by 16.2 percentage points, with effects varying substantially by domain: software engineering gained least, at 4.5 points, and healthcare most, at 51.9 points [14]. SWE-Skills-Bench evaluated 49 public skills across roughly 565 software engineering tasks and found that 39 produced no measurable improvement, with an average gain of 1.2 percent and substantial token overhead [13]. SWE-Skills-Bench attributed poor results in part to *concept bleed*, in which an agent conflates distinct topics presented together in a comprehensive document, and *context-window displacement*, in which skill text crowds out the task itself [13]. SkillsBench v1 similarly reported that focused skills with two or three modules outperformed comprehensive documentation, while self-generated skills provided no benefit on average, suggesting that models do not reliably author the procedural knowledge from which they benefit [14]. Together, these findings suggest that skill effectiveness depends in part on whether a task requires specialized knowledge that the underlying model does not already possess, and on how that knowledge is organized and delivered. Querying a particular KG release is a task of this kind because release-specific identifiers, relationships, source conventions, and graph structures cannot be inferred from the query language alone.

These findings and our own experience motivated much of the DDKG skill’s underlying design. To support complex schemas without overloading LLM context, we designed a small controller with detailed supporting material that was loaded only when needed, and the skill was authored by human experts rather than created by the LLM agent itself. The skill therefore required a modular, programmatic design rather than one large document loaded into the LLM context for every query. Destefanis likewise argues that a skill should be treated as a software artifact, with single responsibility, separation of interface from implementation, low coupling, economy in a shared token budget, and behavioral evaluation [12].

Unlike general Neo4j Agent Skills for Cypher and graph operations [15], ddkg.skill is designed to provide the release-specific knowledge needed to query DDKG, including its identifiers, source conventions, relationships, source-specific structures, validated query patterns, and checks for known failure modes. It helps a general-purpose AI system construct queries that a user can execute against the DDKG, leaving the database as the source of returned results.

### A compositional library for DDKG query generation

The current ddkg.skill release targets the DDKG version DataDistillery_2025_04_DEC. The skill file contains 38 files organized into five parts: a SKILL.md controller, a routing and checking script, 18 curated DDKG reference documents, eight primary DDKG and UBKG documentation sources, and ten structured assets. The structured assets include source and predicate tables, inverse-predicate pairs, endpoint mappings, evidence classes, example identifiers, and the routing table.

The structured assets describe the release broadly rather than only the sources used in worked examples: 289 source abbreviations contribute identifiers, 143 contribute assertions, 2,101 distinct relationship types are represented, and 25,681 subject-predicate-object combinations are classified among the Common Fund datasets. The curated references, by contrast, cover demonstrated query patterns and traps found during testing. This division is deliberate. A source with no prose coverage remains described by the enumerated assets, so questions about datasets that no worked example touches can still be composed from the release inventory.

We call this a compositional library because the agent does not receive one large block of DDKG instructions. Instead, the controller identifies the type of question and directs the agent to the pieces of information needed for that question. The internal file `skill_graph.tsv` links topics to relevant materials using relationships such as `answers, must_read_with, demonstrated_by, invalidated_by,` and `supersedes`. A companion script, `route.py`, checks that these links still point to valid files and sections. This makes it possible to update one rule or source-specific pattern without rewriting a single long prompt.

The curated references are separate from the primary documentation. This distinction matters because some useful DDKG query rules were learned by testing the actual graph rather than by reading the documentation alone. Keeping these layers separate lets the skill record both what the documentation says and what was observed during query testing.

## Design choices

### Five choices determine how the skill behaves

The skill is designed to compose database queries and diagnostic steps but not to connect to the KG directly. The user needs to use the skill results to paste into the Neo4j browser or command line to run each query on their own DDKG instance. Many institutional installations sit behind organizational firewalls where an outbound connection from the data to an external service is not permitted, and this design requires none.

The skill is instructed to not expect any particular query result: no statements around predicted row counts, anticipated entity lists, or claims about what the graph contains. It can specify the columns and row unit needed to make a query interpretable. Each query is accompanied by a check that would show the query went wrong, such as which identifier a correct match should return or what an empty result at a given step would imply.

The skill discussed here is paired to the DDKG December 2025 release. The facts that make queries succeed, including which source abbreviations name relationships rather than record namespaces, how quantitative values are binned, and which member of an inverse pair a traversal can bind, will differ between DDKG releases. An explicit hierarchical instruction in the skill resolves conflicts in the skill’s internal structure: validation executed against the release outranks release documentation, which in turn outranks framework documentation. That order was adopted after the documented names of structural edges connecting concepts to identifiers changed between releases, and queries written from documentation returned nothing while raising no error.

Work is handed over in stages when necessary for more complex queries. The skill writes one step, the user runs it, and the next step is written against the returned values. Where a question requires a scientific judgment, such as which of several similarly named pathway definitions to adopt, the choice returns to the user together with the data that informs the decision.

Finally, the skill is not optimized for a particular AI vendor or product, and the same archive was exercised by both open source as well as commercial assistants during development. It is paired to the DDKG graph release, not to a particular LLM model.

### How ddkg.skill works

The skill uses a simple staged workflow. It first looks for a validated query pattern that is close to the user’s question. It then resolves the main entity, asks for clarification when a term is ambiguous, and prefers stable DDKG identifiers when possible. If the route through the graph is uncertain, the skill generates inspection queries to determine which sources and relationships connect to the entity.

The final Cypher is assembled using DDKG-specific source, predicate, and endpoint information. Fixed rules address common problems such as inverse relationships, species restrictions, missing properties, grouping, row limits, and duplicated graph representations. The skill also asks what one result row is supposed to represent. This prevents graph-level duplication from being mistaken for multiple genes, diseases, compounds, or other biological entities.

The skill passes the database query to the user, who can then run the Cypher against a compatible DDKG instance. Only the returned rows are interpreted. In this way, examples in the skill teach query structure rather than acting as stored answers.

### Biomedical use cases

To illustrate how ddkg.skill supports a researcher starting from a biological question rather than a graph schema, we applied the current skill to three biomedical queries. In each example, the user supplied ddkg.skill and a single natural-language question. The assistant generated Cypher that was then executed against the local December 2025 DDKG instance. These examples demonstrate practical query construction and interpretation; they are separate from the orthogonal nine-test evaluation described below. The exact generated Cypher for all three use cases is provided in Supplementary File S1.

### Use case 1: Cross-species phenotype evidence to adult human heart expression

The first question asked, “Which genes associated with atrial septal defects in mice are expressed in human heart tissues in GTEx?” The generated query anchored the Mammalian Phenotype Ontology term `MP:0010403` and more specific phenotypes within four `isa` levels, followed MPMGI phenotype-to-mouse-gene relationships, mapped mouse genes to human 1:1 orthologs through HCOP, and then queried `GTEXEXP` for nonzero expression in the adult heart atrial appendage and left ventricle. This route preserves the evidence type in the question: the starting evidence is a mouse phenotype rather than a human disease association.

Execution returned 181 human genes with nonzero expression in at least one of the two adult GTEx heart tissues (**Figure 1**). Because GTEx represents adult tissue, low or absent expression does not exclude a gene that acts transiently during atrial septation, but it provides an adult expression filter on model-organism phenotype evidence, rather than a complete test of developmental relevance.

**Figure 1.**
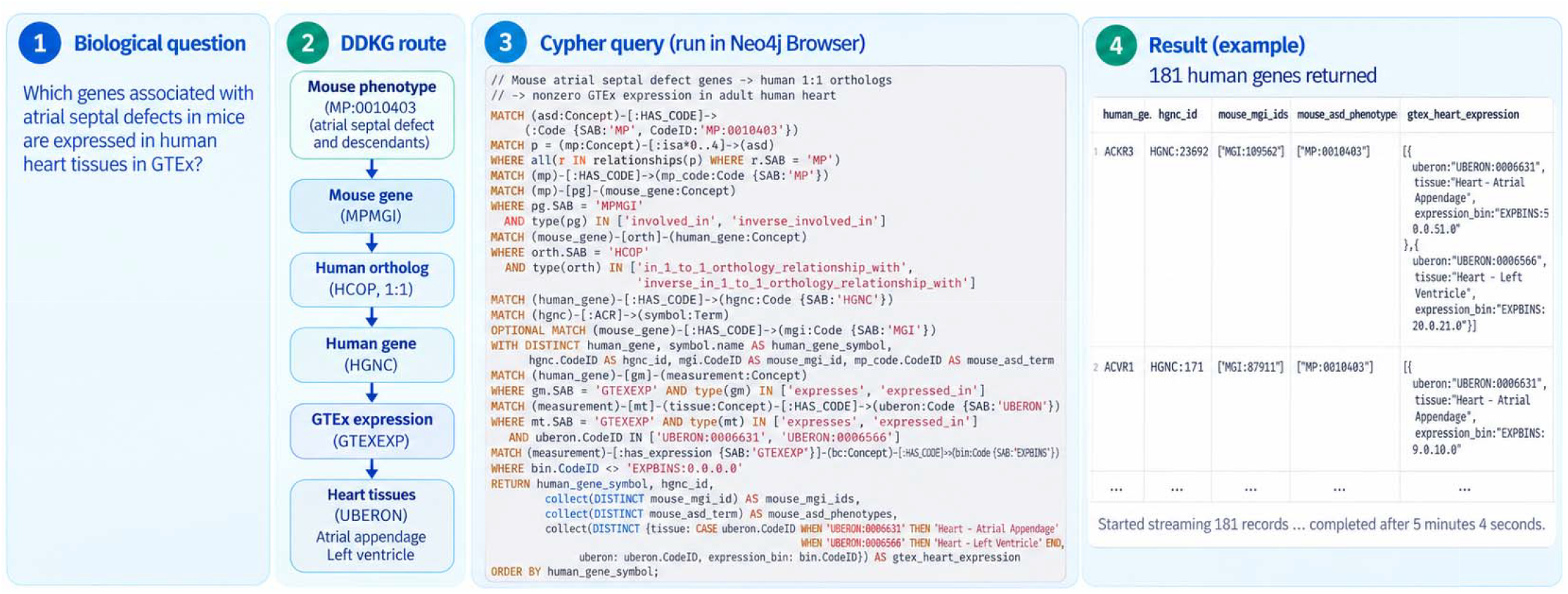
Use case 1: connecting mouse atrial septal defect phenotypes to human orthologs and adult heart expression. Starting from MP:0010403, ddkg.skill generated a traversal through MPMGI mouse phenotype evidence, HCOP 1:1 orthology, HGNC genes, and GTEx expression in heart atrial appendage and left ventricle. The executed query returned 181 human genes with nonzero expression in at least one of the two tissues. The figure shows the biological question, graph route, generated Cypher, and representative Neo4j output.

### Use case 2: GTEx eQTL overlap with 4DN chromatin-loop anchors

The second question asked, “Which GTEx eQTLs overlap 4DN chromatin-loop anchor regions, and which genes are they associated with?” The generated query used the four `4DN` anchor boundary relationships to define upstream and downstream anchor intervals. `HSCLO` 1 kbp bins provided the shared genomic coordinate layer for matching `GTEXEQTL` records to the anchor intervals, after which the query retrie ed the associated `HGNC` genes. Importantly, the overlap was restricted to anchor intervals rather than the full span between loop anchors.

The demonstration intentionally limited the intermediate result to 5,000 rows before adding display fields. In that 5,000-row slice, the output contained 1,378 unique eQTL records, 372 unique genes, 339 unique 4DN loops, and 759 unique 1 kbp `HSCLO` bins (Figure 2). The 5,000 returned rows represented 2,500 unique eQTL-gene rows because reciprocal located_in and location_of relationships were both present. The example therefore also illustrates why row grain and inverse relationships need to be considered when interpreting graph query output.

**Figure 2.**
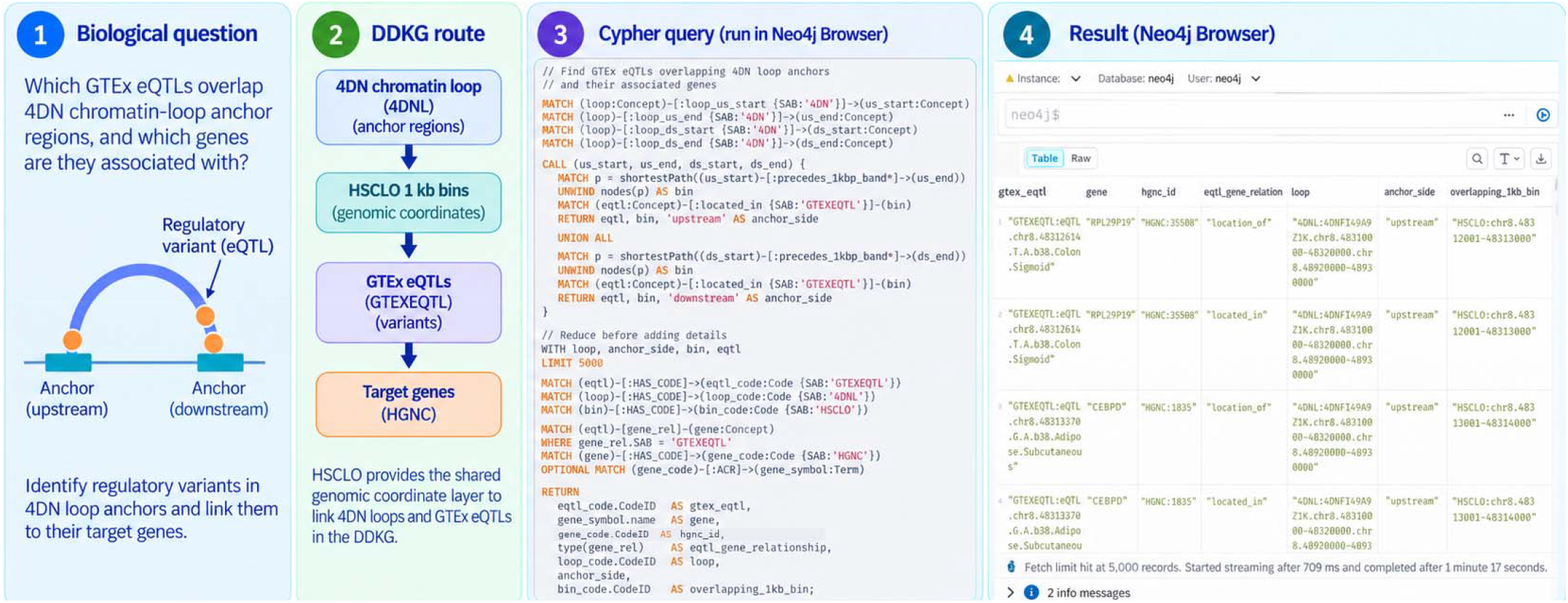
Use case 2: identifying GTEx eQTLs that overlap 4DN chromatin-loop anchor regions. ddkg.skill generated a query that expands the upstream and downstream 4DN anchor intervals through HSCLO 1 kbp bins, matches GTEx eQTL records to those bins, and retrieves their HGNC gene associations. The displayed run is a 5,000-row slice rather than an exhaustive census. It contains 1,378 unique eQTL records, 372 genes, 339 loops, and 759 unique 1 kbp bins. Reciprocal eQTL-gene relationships produce two graph rows for each of 2,500 unique eQTL-gene combinations in the slice.

### Use case 3: Congenital heart phenotype, ClinVar evidence, and heart expression

The third question asked, “Which genes associated with tetralogy of Fallot have pathogenic or likely pathogenic ClinVar variants, and which of those genes are also expressed in human heart tissues in GTEx?” The query anchored Tetralogy of Fallot at HP:0001636, collected the provenance and predicates for directly associated HGNC genes, required a ClinVar gene-condition assertion, tested whether the ClinVar condition was directly Tetralogy of Fallot, and annotated nonzero GTEx expression in adult heart atrial appendage and left ventricle. In this DDKG release, the ClinVar ingestion itself was restricted to pathogenic, likely pathogenic, and pathogenic/likely pathogenic single nucleotide variants, so the generated query uses the presence of CLINVAR assertions rather than a separate clinical-significance property filter.

The executed query returned 131 gene rows and recorded GTEx heart expression as a Boolean together with the expression bins, allowing expressed genes to be separated from genes without nonzero expression in those adult tissues (Figure 3). The displayed rows include GATA6 and JAG1, both with direct ClinVar Tetralogy of Fallot evidence and nonzero GTEx heart expression. This example shows how the skill can preserve source provenance while combining phenotype, clinical variant, and tissue-expression evidence in one result table.

**Figure 3.**
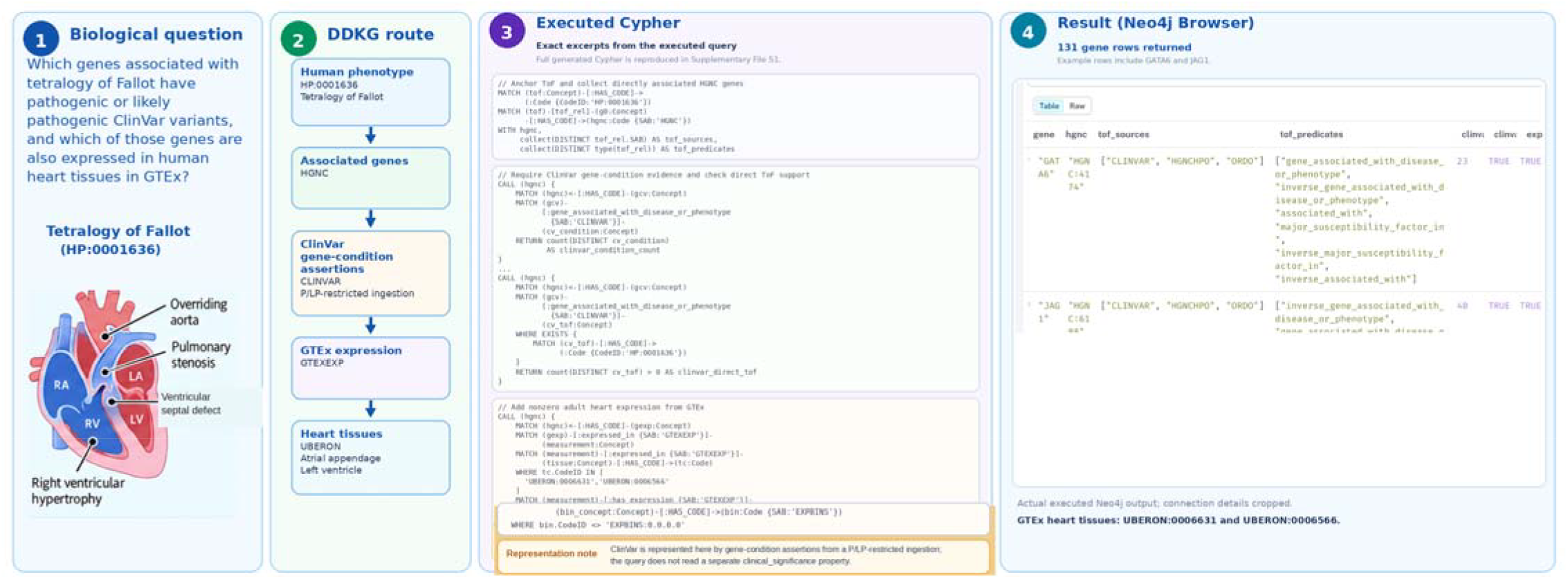
Use case 3: integrating Tetralogy of Fallot gene associations, ClinVar evidence, and GTEx heart expression. The generated query anchors HP:0001636, records the sources and predicates supporting associated genes, requires ClinVar gene-condition evidence, checks whether ClinVar directly links the gene to Tetralogy of Fallot, and adds nonzero adult heart expression from GTEx. The run returned 131 gene rows with expression status and expression-bin metadata. The result table preserves the distinction between general ClinVar condition support and a direct ClinVar Tetralogy of Fallot association.

### Testing and revision

The ddkg.skill package was developed by repeatedly generating Cypher and running it against real DDKG instances on various LLM platforms. This process uncovered errors that were easy to miss by visual inspection alone, including outdated edge names, source-specific relationship directions, incorrect identifier assumptions, duplicated concepts, hidden row multiplication, species mismatches, and graph patterns that require an intermediate association node.

The public repository includes a lightweight behavioral test protocol with regression, novel, ambiguity, evidence, adversarial, scale, and result-unit cases. Tests use fresh conversations, record the generated Cypher, and judge success from actual execution. Older test results are retained as examples of the development process but are marked as superseded when they do not apply to the current skill archive. The orthogonal nine-test series reported here was run on an earlier version of the ddkg.skill: the R5b-rebased build (297,218 bytes, 38 files, 227 routing entries against DataDistillery_2025_04_DEC on Neo4j 5.26.28 Community. Supplementary File S1 summarizes the test design, execution, and outcomes. The public repository preserves the complete historical Tier 7 record, including the recovered original query interactions and generated Cypher.

Failures are sorted into four categories that require different repairs: a reference states something false; the relevant fact was present but not consulted; the fact is absent; or the answer is correct but unhelpful. The second category dominated early testing and led directly to the routing table, since a reader who knows where a subject is documented may still not know what else bears on it. The most consequential category was the first. Statements that were false and produced silently empty results were found only by executing queries and reading the output, and two such statements survived several rounds of review before a test exposed them.

We designed a nine-question series around problems an earlier version of the skill had not yet seen in the worked examples used to build it. These included: 1) which genes are associated with Ehlers-Danlos syndrome in ClinVar but have no reported eQTL in GTEx; 2) what DisGeNET reports for hypertrophic cardiomyopathy; 3) whether PCSK9 has strongly co-expressed genes in GTEx, 4) which genes exceed 100 TPM expression in pancreas, 5) which mouse phenotypes are linked to human orthologs of ciliary transition-zone genes, 6) whether ClinGen gene-disease assertions could meaningfully be compared with Open Targets Genetics, 7) which kidney marker genes are represented in Azimuth; 8) which drugs MED-RT lists as contraindicated for long QT syndrome and 9) separating the answerable and unanswerable parts of a question about electrophysiological properties of epilepsy-associated ion channels. The overview and results of these queries are presented in the supplementary file S1.

The value of this test series was not simply the number of queries that succeeded. Five of the seven questions involving previously unused data sources produced correct executed results in the version of the ddkg.skill we tested. More importantly, the failures showed what the skill could and could not recognize. For the ClinGen and Open Targets Genetics question, it correctly determined that the requested gene-level comparison was not meaningful because the two sources connect to different entity types in this DDKG release. The pancreas expression query succeeded only after execution exposed two incorrect assumptions in the skill’s own reference material. The ion-channel question was correctly divided into a part the DDKG could answer and electrophysiological measurements that it does not contain.

A methodological observation from this process revealed an important design choice. Rules or positive statements that can be checked directly in the generated query were applied more consistently than cautious or negative statements that depend on the agent recognizing, during composition, that an adversarial situation applied. For instance, an early caution given to the skill about ordering a result by a column with few distinct values, and then truncating it, followed this pattern. For the affected query pattern, we changed it so that it was restated as a countable diagnostic: if the ordering column has fewer distinct values than the row limit, truncation may cut through tied groups and the query should be inspected. Cautions that could not be reduced to such a condition were moved into the checking script, where they run whether the agent remembers them or not.

## Discussion

The main idea behind ddkg.skill is simple: a complex scientific graph needs a body of practical, version-specific knowledge that can be maintained alongside the graph. The Agent Skills format provides a convenient way to package that knowledge without changing model weights. The compositional library keeps the main instructions small while allowing the agent to retrieve detailed DDKG rules only when they are relevant.

The compositional structure also speaks to reported limits of skills in other domains. Concept bleed and context window displacement, the two failure modes identified in software engineering evaluations [13], arise when an agent is given a large body of information that includes material unrelated to the immediate task. Separating a small controller from material loaded only when a routing entry calls for it keeps the amount of skill text in context small and keeps unrelated topics out at the same time. The design is consistent with the observation that focused skills outperform comprehensive documentation [14], although we have not measured module count against performance directly.

Two limitations bound these claims. Whether the approach generalizes beyond one graph is untested. The evaluation reported here is behavioral, judging whether composed queries run and return interpretable results, rather than a controlled comparison against the same assistant without the skill. The paired design used by both published benchmarks [13,14] would give a quantitative estimate of benefit for this domain, and applying it to a biomedical knowledge graph is a natural next study.

This approach also makes planned graph changes easier to handle. The December 2025 skill is tied to that DDKG schema. A future DDKG release based on the JSON Knowledge Graph representation will require a different skill version rather than assuming that old query rules still apply. The same principle may be useful for other scientific databases in which schemas, identifiers, and source-specific meaning change over time.

## Software availability

The DDKG Agent Skill, installation instructions, and behavioral testing materials are available at https://github.com/TaylorResearchLab/ddkg-agent. The current R7 skill targets the December 2025 DDKG release (DataDistillery_2025_04_DEC) and is distributed as a single archive of 38 files (303,258 bytes; SHA-256 eca6c7f7461bab113160fe8a4c71260c35fc4350a0b632294d56ef770cbfdb60). Supplementary File S1 summarizes the orthogonal nine-test evaluation and provides the exact Cypher generated for the three demonstration use cases. The complete historical Tier 7 evaluation record, including recovered original generated Cypher, execution status, returned results, and revisions, is maintained at https://github.com/TaylorResearchLab/ddkg-agent/blob/main/ddkg-agent-skill/ddkg_tier7_complete_record.md.

