## Supplemental File S1 for "ddkg.skill: A Compositional Agent Skill for Translating Biomedical and Bioinformatics Questions into Cypher for the Data Distillery Knowledge Graph"

Supplementary File S1

Orthogonal nine-test evaluation of ddkg.skill

| **Evaluation build** | R5b-rebased build; 38 files; 227 routing relationships; SHA-256 c092480a61f2e46efb3586aacfe2240a3cebb83a0c769c5dfa0eb5ea54b8c52b |
| --- | --- |
| **DDKG release** | DataDistillery_2025_04_DEC (December 2025 release) |
| **Database software** | Neo4j 5.26.28 Community |
| **Execution dates** | 2 September 2026 and 8 September 2026 |
| **Purpose** | Evaluate whether ddkg.skill could construct useful DDKG queries for data sources and query patterns that were not represented in its worked examples. |
| **Relation to current skill** | The manuscript describes a later R7 skill build with 38 files and 239 routing relationships. The nine tests below were performed on the earlier R5b-rebased build. Findings from these tests informed later revisions; the tests were not rerun on R7 for this report. |

**How to read this supplement.** Seven tests asked about DDKG sources that were not represented in the skill's worked examples. Two tests focused instead on query constructions that were not represented in those examples. Each test was judged from queries actually executed against the December 2025 DDKG release, not from whether the generated Cypher merely looked plausible. A "pass after correction" means that an initial problem was identified during execution, the query or skill reference material was corrected, and the revised query then met the test criteria. A "partial pass" means that part of the question was handled correctly but the answerable query did not complete successfully.

Complete historical record. This supplement is a concise summary of the orthogonal evaluation. The complete Tier 7 record, including recovered original query interactions, generated Cypher, execution status, returned results, and revisions, is preserved in the public repository at https://github.com/TaylorResearchLab/ddkg-agent/blob/main/ddkg-agent-skill/ddkg_tier7_complete_record.md. Recovered queries are retained there exactly as generated, including failed and superseded attempts where they were part of the evaluation history.

### Overall results

| **Test** | **Question area** | **Outcome** |
| --- | --- | --- |
| T7.1 | ClinVar plus GTEx eQTL | Fail |
| T7.2 | DisGeNET | Pass |
| T7.3 | GTEx co-expression | Pass |
| T7.4 | GTEx expression threshold | Pass after correction |
| T7.5 | Human-to-mouse orthology and mouse phenotypes | Pass with one corrected conclusion |
| T7.6 | ClinGen versus Open Targets Genetics | Requested comparison not valid at the gene level |
| T7.7 | Azimuth kidney markers | Pass; species caveat omitted |
| T7.8 | MED-RT contraindications | Pass |
| T7.9 | Epilepsy ion-channel electrophysiology | Partial pass |

### T7.1. Which genes are associated with Ehlers-Danlos syndrome in ClinVar but have no reported eQTL in GTEx?

| **What this test asked** | Use ClinVar and GTEx eQTL data to identify ClinVar-associated genes and then exclude genes represented in the GTEx eQTL set. |
| --- | --- |
| **What counted as success** | Resolve Ehlers-Danlos syndrome without dropping relevant vocabularies; identify ClinVar-associated genes; exclude genes present in the GTEx eQTL set; and state that the GTEx eQTL data in this DDKG build are a filtered subset rather than a complete record of all GTEx eQTLs. |
| **What happened** | The first query returned 100 rows, but the row limit was reached before all relevant vocabularies were represented. The next query did not complete. |
| **Outcome** | Fail. |
| **Interpretation** | The disease-resolution step was incomplete, and the second query was too broad and computationally expensive to finish. The test therefore did not produce a valid gene list. |

### T7.2. What does DisGeNET say about hypertrophic cardiomyopathy?

| **What this test asked** | Use DisGeNET, a large DDKG source that was not represented in the worked examples. |
| --- | --- |
| **What counted as success** | Resolve the disease; use the DisGeNET relationships present in the graph; and preserve the different DisGeNET association classes rather than treating every record as the same type of causal evidence. |
| **What happened** | The disease resolved to a concept with 5,569 DisGeNET partners. The returned records belonged to several association classes, including variant-disease, gene-variant, biomarker, causal-mutation, age-of-onset, gene-disease, and therapeutic classes. |
| **Outcome** | Pass. |
| **Interpretation** | The executed queries showed that DisGeNET is represented by several distinct kinds of associations. Results should therefore be reported by association class rather than as one undifferentiated count. |

### T7.3. Which genes are strongly co-expressed with PCSK9?

| **What this test asked** | Use the GTEXCOEXP source, which was not represented in the worked examples. |
| --- | --- |
| **What counted as success** | Recognize that a GTEXCOEXP edge already means that the gene pair met the source criterion of Pearson correlation above 0.99 in at least one tissue. Do not invent a stored correlation coefficient or impose a new numeric threshold. |
| **What happened** | The initial co-expression query returned no rows. The assistant then checked the structural schema, resolved PCSK9 to HGNC:20001, profiled all relationships touching the PCSK9 concept, and finally queried GTEXCOEXP directly. That source-specific check also returned no rows, establishing the empty result as a true negative for this DDKG representation rather than a schema or entity-resolution failure. Additional STRING and MSigDB queries were then used only as biologically distinct alternatives, not as substitutes for GTEx co-expression. |
| **Outcome** | Pass. |
| **Interpretation** | An empty result can be a valid database result. The skill used the source as it is represented in the DDKG and did not invent a ranking score when no partners were returned. |

### T7.4. List genes expressed above 100 TPM in pancreas.

| **What this test asked** | Test a numeric expression threshold using GTEXEXP. This threshold-style query pattern was not represented in the worked examples. |
| --- | --- |
| **What counted as success** | Use the expression-bin identifiers actually stored in the graph rather than assuming that expression is a numeric node property; verify how the bins are encoded before applying the threshold; and return one row per gene. |
| **What happened** | The first attempts failed because the documented expression-bin format and one assumed tissue-side relationship were incorrect. Diagnostic queries identified the pancreas anchor, the relationship used on the tissue side, and the bin identifiers actually present in the graph. The final query returned 314 distinct HGNC genes. |
| **Outcome** | Pass after correction. |
| **Interpretation** | Execution exposed two incorrect statements in the skill reference material. After those assumptions were corrected, the query used graph-observed bin identifiers and returned a nonduplicated gene list. These findings were incorporated into later skill revisions. |

### T7.5. Which mouse phenotypes are associated with human orthologs of genes in the ciliary transition zone?

| **What this test asked** | Test HCOP human-to-mouse orthology, MPMGI mouse phenotype data, and construction of a ciliary-transition-zone gene set. |
| --- | --- |
| **What counted as success** | Use the human-to-mouse orthology source and the mouse phenotype source; identify returned phenotypes as mouse phenotypes; and do not treat the ciliary transition zone as though it were itself a single gene. |
| **What happened** | A profile of GO:0035869 found no direct HGNC gene links, but it found 67 UniProtKB proteins connected to the term, providing a possible graph-based route to genes. The formal Tier 7 cross-species query used a curated 17-gene input set and returned mouse phenotype data for 16 of 17 genes; TMEM216 was the only seed absent from that phenotype output. A separate follow-up query later showed that TMEM216 did have a 1:1 HCOP mouse ortholog but no MPMGI phenotype assertions in this DDKG release. |
| **Outcome** | Pass with one corrected conclusion. |
| **Interpretation** | The cross-species HCOP-to-MPMGI route worked and the returned phenotypes were correctly identified as mouse mutant/knockout phenotype evidence. The later TMEM216 diagnostic shows that its absence from the formal output reflected an MPMGI annotation or load gap rather than failure of the HCOP orthology mapping. An initial conclusion that the transition-zone gene set had to come entirely from outside the graph was also too strong because a UniProtKB protein-layer route to GO:0035869 exists. |

### T7.6. Which genes have ClinGen gene-disease validity assertions but no Open Targets Genetics variant-disease association?

| **What this test asked** | Test ClinGen and Open Targets Genetics, neither of which was represented in the worked examples, and determine whether a gene-level set subtraction between the two sources is meaningful in this graph. |
| --- | --- |
| **What counted as success** | Before subtracting one source from the other, verify that both sources connect at the same entity level. Return ClinGen classification values and do not interpret absence from Open Targets Genetics as biological evidence. |
| **What happened** | The Open Targets Genetics relationship connected genomic locations and disease concepts rather than HGNC genes. Among 1,935 ClinGen genes, zero could connect directly to the Open Targets Genetics relationship at the gene level. |
| **Outcome** | The requested gene-level comparison is not valid. |
| **Interpretation** | The two sources do not meet at the gene level in this DDKG release, so a gene list created by subtracting one source from the other would be misleading. The skill warned that this connection needed to be checked, although it also composed the downstream subtraction query. This test illustrates why diagnostic steps should be executed before continuing to a cross-source comparison. |

### T7.7. What cell-type marker genes does Azimuth provide for kidney?

| **What this test asked** | Test Azimuth kidney marker data, a small source that was not represented in the worked examples. |
| --- | --- |
| **What counted as success** | Use the Azimuth marker relationship; check the source size so that a row limit does not silently truncate the result; and note that Azimuth reference atlases can include more than one species. |
| **What happened** | The executed query returned 49 kidney cell types and 485 marker assignments, exactly matching the total number of kidney marker relationships recorded for the source. |
| **Outcome** | Pass for the query and result; one caveat was omitted. |
| **Interpretation** | The source was recovered correctly, including the distinction between AZ nodes and HMAZ marker relationships. This distinction corrected an assumption in the original test description, which had conflated those two source abbreviations. The explanation still omitted the required species check. |

### T7.8. Which drugs are contraindicated in patients with long QT syndrome?

| **What this test asked** | Test MED-RT contraindication relationships. The source had no prose coverage in the skill and therefore tested whether the structured source inventory could support query construction. |
| --- | --- |
| **What counted as success** | Find the MED-RT contraindication relationships from the machine-readable source inventory; distinguish MED-RT labeling statements from an independent clinical risk classification; and retain class-level results if the graph contains them. |
| **What happened** | The query returned 17 distinct concept rows, including disopyramide, dronedarone, nilotinib, and sotalol. For this disease anchor, only the individual-drug contraindication relationship returned rows. |
| **Outcome** | Pass. |
| **Interpretation** | The skill found a contraindication source that the original test design had assumed was absent from the graph. It recovered the relevant MED-RT relationship from the structured registry even though that source was not described in the prose references. Some display labels were poor synonyms, but the underlying concept rows were correct. |

### T7.9. What are the electrophysiological parameters for ion channels associated with epilepsy?

| **What this test asked** | Test a question with one part that the DDKG can address and another part that it cannot. This was a query-construction test rather than a test of a previously uncovered source. |
| --- | --- |
| **What counted as success** | Separate the question into two parts. State that channel conductance, gating, kinetics, voltage dependence, and permeability are not represented in this DDKG release. Do not invent relationships for those measurements. Attempt the answerable gene-association portion separately. |
| **What happened** | The skill correctly stated that the requested electrophysiological measurements are not represented and named external resources where such measurements are typically found. The disease-resolution query returned a mixture of human and nonhuman terms. The next gene query crashed the database instance, and a similar follow-up query was not run. |
| **Outcome** | Partial pass. |
| **Interpretation** | The unavailable electrophysiology portion was handled correctly. The answerable gene-association portion was not completed successfully, so the full test did not pass. |

### Interpretation

This evaluation was performed on the R5b-rebased build of ddkg.skill. The results show that the skill could use structured source and relationship inventories beyond its worked examples, but they also show why execution against the actual graph remains necessary. The tests exposed incomplete term resolution, source representations that did not connect at the entity level requested by the user, expensive queries that did not finish, and incorrect assumptions in the skill reference material itself. Several findings directly informed later revisions of the skill. The public Tier 7 historical record preserves the recovered original query interactions and generated Cypher so that the summarized outcomes here can be traced back to the actual query-construction sequence. The evaluation should therefore be read as a record of both successful behavior and failure modes, rather than as evidence that every generated query will be correct without execution and checking.

### Demonstration use cases and generated Cypher queries

The following three use cases are separate from the orthogonal nine-test evaluation above. In each case, the user supplied ddkg.skill and a single biomedical question. The assistant generated Cypher, and the user executed that query against the local December 2025 DDKG instance. The queries are reproduced here as generated so that readers can see the exact graph traversal used. These examples demonstrate workflow and should not be interpreted as a controlled benchmark of the skill.

#### Use case 1: Mouse atrial septal defect evidence to human heart expression

**User question.** Which genes associated with atrial septal defects in mice are expressed in human heart tissues in GTEx?

**Purpose.** Follow mouse atrial septal defect phenotype evidence through mouse genes and human 1:1 orthologs, then retain human genes with nonzero adult heart expression in GTEx atrial appendage or left ventricle.

**Observed result.** The Neo4j execution returned 181 human genes. The displayed example rows include ACKR3 and ACVR1 with mouse MGI identifiers, MP:0010403 phenotype evidence, and GTEx heart expression bins. The query searches MP:0010403 and descendants up to four isa steps, so the ontology expansion is bounded rather than unlimited. GTEx is an adult-tissue filter and should not be used to exclude genes with transient roles in heart development.

##### Generated Cypher

```
// Mouse atrial septal defect genes -> human 1:1 orthologs
// -> nonzero GTEx expression in adult human heart

// 1. Anchor mouse atrial septal defect
MATCH (asd:Concept)-[:HAS_CODE]->
      (:Code {SAB:'MP', CodeID:'MP:0010403'})

// Include the ASD term itself and more specific ASD phenotypes.
// MP isa is child -> parent.
MATCH p = (mp:Concept)-[:isa*0..4]->(asd)
WHERE all(r IN relationships(p) WHERE r.SAB = 'MP')

MATCH (mp)-[:HAS_CODE]->(mp_code:Code {SAB:'MP'})

// 2. Mouse phenotype -> mouse gene
MATCH (mp)-[pg]-(mouse_gene:Concept)
WHERE pg.SAB = 'MPMGI'
  AND type(pg) IN ['involved_in', 'inverse_involved_in']

WITH DISTINCT mouse_gene, mp_code

// 3. Mouse gene -> human 1:1 ortholog
MATCH (mouse_gene)-[orth]-(human_gene:Concept)
WHERE orth.SAB = 'HCOP'
  AND type(orth) IN [
    'in_1_to_1_orthology_relationship_with',
    'inverse_in_1_to_1_orthology_relationship_with'
  ]

MATCH (human_gene)-[:HAS_CODE]->(hgnc:Code {SAB:'HGNC'})
MATCH (hgnc)-[:ACR]->(symbol:Term)

OPTIONAL MATCH (mouse_gene)-[:HAS_CODE]->(mgi:Code {SAB:'MGI'})

WITH DISTINCT
     human_gene,
     symbol.name AS human_gene_symbol,
     hgnc.CodeID AS hgnc_id,
     mgi.CodeID AS mouse_mgi_id,
     mp_code.CodeID AS mouse_asd_term

// 4. Human ortholog -> GTEx measurement
MATCH (human_gene)-[gm]-(measurement:Concept)
WHERE gm.SAB = 'GTEXEXP'
  AND type(gm) IN ['expresses', 'expressed_in']

// 5. Restrict to the two GTEx adult heart tissues
MATCH (measurement)-[mt]-(tissue:Concept)
      -[:HAS_CODE]->(uberon:Code {SAB:'UBERON'})
WHERE mt.SAB = 'GTEXEXP'
  AND type(mt) IN ['expresses', 'expressed_in']
  AND uberon.CodeID IN [
      'UBERON:0006631',   // Heart - Atrial Appendage
      'UBERON:0006566'    // Heart - Left Ventricle
  ]

// 6. Require actual nonzero expression.
// A GTEXEXP measurement alone does not imply TPM > 0.
MATCH (measurement)-[:has_expression {SAB:'GTEXEXP'}]-
      (bc:Concept)-[:HAS_CODE]->(bin:Code {SAB:'EXPBINS'})
WHERE bin.CodeID <> 'EXPBINS:0.0.0.0'

RETURN
    human_gene_symbol,
    hgnc_id,
    collect(DISTINCT mouse_mgi_id) AS mouse_mgi_ids,
    collect(DISTINCT mouse_asd_term) AS mouse_asd_phenotypes,
    collect(DISTINCT {
        tissue:
            CASE uberon.CodeID
                WHEN 'UBERON:0006631'
                    THEN 'Heart - Atrial Appendage'
                WHEN 'UBERON:0006566'
                    THEN 'Heart - Left Ventricle'
            END,
        uberon: uberon.CodeID,
        expression_bin: bin.CodeID
    }) AS gtex_heart_expression
ORDER BY human_gene_symbol;
```

#### Use case 2: 4DN chromatin-loop anchors and GTEx eQTLs

**User question.** Which GTEx eQTLs overlap 4DN chromatin-loop anchor regions, and which genes are they associated with?

**Purpose.** Define each 4DN upstream and downstream anchor using its start and end HSCLO bins, identify GTEx eQTL records that occupy the same 1 kbp bins, and return the genes associated with those eQTL records.

**Observed result.** The query intentionally limits the intermediate output to 5,000 rows. The displayed slice contained 1,378 unique eQTL records, 372 unique genes, 339 unique 4DN loops, and 759 unique 1 kbp HSCLO bins. Those 5,000 rows represented 2,500 unique eQTL-gene combinations because both located_in and location_of relationship directions were returned. The result is therefore a demonstration slice, not an exhaustive count of all overlaps in DDKG.

##### Generated Cypher

```
MATCH (loop:Concept)-[:loop_us_start {SAB:'4DN'}]->(us_start:Concept)
MATCH (loop)-[:loop_us_end {SAB:'4DN'}]->(us_end:Concept)
MATCH (loop)-[:loop_ds_start {SAB:'4DN'}]->(ds_start:Concept)
MATCH (loop)-[:loop_ds_end {SAB:'4DN'}]->(ds_end:Concept)

CALL (us_start, us_end, ds_start, ds_end) {

    MATCH p = shortestPath(
        (us_start)-[:precedes_1kbp_band*]->(us_end)
    )
    UNWIND nodes(p) AS bin
    MATCH (eqtl:Concept)-[:located_in {SAB:'GTEXEQTL'}]-(bin)
    RETURN eqtl, bin, 'upstream' AS anchor_side

    UNION ALL

    MATCH p = shortestPath(
        (ds_start)-[:precedes_1kbp_band*]->(ds_end)
    )
    UNWIND nodes(p) AS bin
    MATCH (eqtl:Concept)-[:located_in {SAB:'GTEXEQTL'}]-(bin)
    RETURN eqtl, bin, 'downstream' AS anchor_side
}

// CRITICAL: reduce here, before decorating the rows
WITH loop, anchor_side, bin, eqtl
LIMIT 5000

MATCH (eqtl)-[:HAS_CODE]->(eqtl_code:Code {SAB:'GTEXEQTL'})
MATCH (loop)-[:HAS_CODE]->(loop_code:Code {SAB:'4DNL'})
MATCH (bin)-[:HAS_CODE]->(bin_code:Code {SAB:'HSCLO'})

MATCH (eqtl)-[gene_rel]-(gene:Concept)
WHERE gene_rel.SAB = 'GTEXEQTL'

MATCH (gene)-[:HAS_CODE]->(gene_code:Code {SAB:'HGNC'})
OPTIONAL MATCH (gene_code)-[:ACR]->(gene_symbol:Term)

RETURN
    eqtl_code.CodeID AS gtex_eqtl,
    gene_symbol.name AS gene,
    gene_code.CodeID AS hgnc_id,
    type(gene_rel) AS eqtl_gene_relationship,
    loop_code.CodeID AS loop,
    anchor_side,
    bin_code.CodeID AS overlapping_1kb_bin;
```

#### Use case 3: Tetralogy of Fallot, ClinVar, and GTEx heart expression

**User question.** Which genes associated with tetralogy of Fallot have pathogenic or likely pathogenic ClinVar variants, and which of those genes are also expressed in human heart tissues in GTEx?

**Purpose.** Retrieve genes directly associated with Tetralogy of Fallot, require ClinVar gene-condition evidence, distinguish a direct ClinVar Tetralogy of Fallot association from other ClinVar conditions, and annotate nonzero adult heart expression in GTEx.

**Important representation note.** The DDKG ClinVar ingestion used for this release was already restricted to pathogenic, likely pathogenic, and pathogenic/likely pathogenic single nucleotide variants. The generated query therefore does not read a separate clinical_significance property. Instead, presence of a CLINVAR gene-condition assertion is the evidence gate. The query also annotates GTEx heart expression but does not remove rows with no adult heart expression; the expressed_in_gtex_heart Boolean is the field used to make that distinction.

**Observed result.** The execution returned 131 gene rows. In the displayed examples, GATA6 and JAG1 had ClinVar condition counts of 23 and 40, respectively. Both were marked as having a direct ClinVar Tetralogy of Fallot association and nonzero GTEx adult heart expression.

##### Generated Cypher

```
// Tetralogy of Fallot
// HP:0001636

MATCH (tof:Concept)-[:HAS_CODE]->(:Code {CodeID:'HP:0001636'})

// Direct gene associations with Tetralogy of Fallot
MATCH (tof)-[tof_rel]-(g0:Concept)-[:HAS_CODE]->(hgnc:Code {SAB:'HGNC'})

WITH hgnc,
     collect(DISTINCT tof_rel.SAB) AS tof_sources,
     collect(DISTINCT type(tof_rel)) AS tof_predicates


// ------------------------------------------------------------
// Require pathogenic / likely pathogenic ClinVar evidence
// ------------------------------------------------------------

CALL (hgnc) {

    MATCH (hgnc)<-[:HAS_CODE]-(gcv:Concept)

    MATCH (gcv)-
          [:gene_associated_with_disease_or_phenotype {SAB:'CLINVAR'}]-
          (cv_condition:Concept)

    RETURN count(DISTINCT cv_condition) AS clinvar_condition_count
}

WITH hgnc,
     tof_sources,
     tof_predicates,
     clinvar_condition_count

WHERE clinvar_condition_count > 0


// ------------------------------------------------------------
// Is the ClinVar association specifically to ToF?
// ------------------------------------------------------------

CALL (hgnc) {

    MATCH (hgnc)<-[:HAS_CODE]-(gcv:Concept)

    MATCH (gcv)-
          [:gene_associated_with_disease_or_phenotype {SAB:'CLINVAR'}]-
          (cv_tof:Concept)

    WHERE EXISTS {
        MATCH (cv_tof)-[:HAS_CODE]->(:Code {CodeID:'HP:0001636'})
    }

    RETURN count(DISTINCT cv_tof) > 0 AS clinvar_direct_tof
}

WITH hgnc,
     tof_sources,
     tof_predicates,
     clinvar_condition_count,
     clinvar_direct_tof


// ------------------------------------------------------------
// GTEx adult human heart expression
//
// UBERON:0006631 = right atrial appendage
// UBERON:0006566 = left ventricular myocardium
// ------------------------------------------------------------

CALL (hgnc) {

    MATCH (hgnc)<-[:HAS_CODE]-(gexp:Concept)

    MATCH (gexp)-
          [:expressed_in {SAB:'GTEXEXP'}]-
          (measurement:Concept)

    WHERE measurement <> gexp

    MATCH (measurement)-
          [:expressed_in {SAB:'GTEXEXP'}]-
          (tissue:Concept)

    WHERE tissue <> gexp

    MATCH (tissue)-[:HAS_CODE]->(tc:Code)

    WHERE tc.CodeID IN [
        'UBERON:0006631',
        'UBERON:0006566'
    ]

    MATCH (measurement)-
          [:has_expression {SAB:'GTEXEXP'}]-
          (bin_concept:Concept)-
          [:HAS_CODE]->
          (bin:Code {SAB:'EXPBINS'})

    WHERE bin.CodeID <> 'EXPBINS:0.0.0.0'

    WITH DISTINCT measurement, tc, bin

    RETURN
        count(DISTINCT measurement) AS gtex_heart_measurement_count,
        collect(
            DISTINCT {
                tissue: tc.CodeID,
                expression_bin: bin.CodeID
            }
        ) AS gtex_heart_expression
}

WITH hgnc,
     tof_sources,
     tof_predicates,
     clinvar_condition_count,
     clinvar_direct_tof,
     gtex_heart_measurement_count,
     gtex_heart_expression


// ------------------------------------------------------------
// HGNC symbol
// ------------------------------------------------------------

OPTIONAL MATCH (hgnc)-[:ACR]->(symbol:Term)

WITH
    hgnc,
    tof_sources,
    tof_predicates,
    clinvar_condition_count,
    clinvar_direct_tof,
    gtex_heart_measurement_count,
    gtex_heart_expression,
    head(collect(DISTINCT trim(symbol.name))) AS gene_symbol

RETURN
    gene_symbol,
    hgnc.CodeID AS hgnc_id,
    tof_sources,
    tof_predicates,
    clinvar_condition_count,
    clinvar_direct_tof,
    gtex_heart_measurement_count > 0 AS expressed_in_gtex_heart,
    gtex_heart_expression

ORDER BY
    expressed_in_gtex_heart DESC,
    clinvar_direct_tof DESC,
    gene_symbol

LIMIT 1000;
```
